# Comprehensive RNA-Seq Reveals Molecular Changes in Kidney Malignancy Among People Living with HIV

**DOI:** 10.1101/2021.11.02.466880

**Authors:** Juan Bao, Jianqing Ye, Jingjing Xu, Shanshan Liu, Lin Wang, Zehuan Li, Qiuyue Li, Feng Liu, Xiaomeng He, Heng Zou, Yanling Feng, Christopher Corpe, Xiaoyan Zhang, Jianqing Xu, Tongyu Zhu, Jin Wang

**Author notes:** **Correspondence should be addressed to:** Jin Wang, Ph.D. Shanghai Public Health Clinical Center, Fudan University, 2901 Caolang Road, Jinshan District, Shanghai 201508, China. These authors contributed equally to this work (Juan Bao, Jianqing Ye, Jingjing Xu).

## Abstract

**Background:** Malignancy of the kidney is a rapidly progressive kidney disease and a major source of morbidity and mortality among people living with HIV (PLWH). Patients with HIV-associated kidey cancer experience higher cancer-specific mortality than the general population, and its mechanism remains poorly understood.

**Methods:** To heighten the awareness of kidney malignancy in patients with HIV infection to facilitate the early diagnosis of kidney cancer, we identified 2460 protein-coding transcripts in HIV-associated kidney cancer using comprehensive RNA sequencing (RNA-seq).

**Results:** KISS1R, CAIX, and NPTX2 mRNA expression levels were specifically increased in HIV-associated kidney cancer, and UMOD and TMEM213 mRNA were decreased in most cases based on real-time PCR analyses. These findings were similar to those noted for the general population with renal cell carcinoma. Immunohistochemical staining analysis also showed that a total of 16 of 18 kidney malignant cases among PLWH exhibited positive staining for KISS1R and CAIX.

**Conclusion:** Pathway analysis of the differentially expressed mRNAs in HIV-associated kidney cancer revealed that several key pathways were involved, including voltage-gated chloride channel activity, distal tubule development, collecting duct development, fructose metabolic processes, and negative regulation of lipase activity. The identified molecular changes in kidney malignancy may offer a helpful explanation for cancer progression and open up new therapeutic avenues that may decrease mortality after a cancer diagnosis among PLWH.

## Introduction

Since the introduction of highly active antiretroviral therapy (HAART) in the last two decades, human immunodeficiency virus (HIV)-infected individuals have a significantly reduced incidence of AIDS-related morbidity at the cost of a higher risk for developing certain cancers, and individuals with non-AIDS-defining cancers (NADCs) (1–3) have a worse prognosis than the general population (4). Moreover, non-AIDS-defining cancer has already become one of the top causes of non-AIDS-related death among patients living with HIV (5, 6).

Kidney disease is the fourth-leading cause of death in people living with HIV (PLWH) in the United States and has also become an increasingly important cause of patient morbidity and mortality (7, 8). The native kidney is a reservoir for HIV-1 and can maintain the ability of the virus to persist and produce viral antigens and/or new virions involved in HIV pathogenesis (9). The presence of HIV in kidney cells can manifest itself in various ways, including indolent nephropathy, inflammation, and proteinuria with glomerular abnormalities (10). HIV-positive patients are at increased risk for kidney diseases, such as HIV-associated nephropathy (HIVAN), noncollapsing focal segmental glomerulosclerosis, immune-complex kidney disease, comorbid kidney disease, and kidney injury resulting from prolonged exposure to antiretroviral therapy (11). Zhang recently found that the clinical characteristics, treatment measures and pathology of 19 HIV-positive renal cell carcinoma (RCC) patients were similar to the general population (12). Although similar to renal cell carcinoma, HIV-associated kidney cancer has poor prognosis in an advanced stage and is difficult to detect early due to the lack of molecular biomarkers (13), and the incidence of RCC in the HIV-positive population was greater than that in the non-HIV population (14, 15). In a large meta-analysis of seven population-based HIV cancer studies involving more than 400,000 HIV-positive patients, Grulich reported a standardized incidence ratio of 1.50 for RCC in the HIV-positive population (14).

Although no clear relationship between the degree of immunosuppression and the risk of kidney cancer has been defined, genetic alterations along with dysregulation of epigenetic pathways of HIV-associated kidney cancer are involved in its tumorigenesis as a heterogeneous disease (16). Genomic analysis yielded millions of protein-coding transcripts and noncoding RNAs that participate in virtually all cancer cellular processes, demonstrating that a spectrum of diverse genomic alterations can define renal carcinoma subtypes in the general population with renal cell carcinoma (17). To improve treatment, provide information about the course of cancer, and predict response to chemotherapy, genome-wide expression studies may provide an unbiased approach for investigating the mechanisms of kidney carcinogenesis in HIV infection and molecularly defining this cancer. In this report, we employed a comprehensive RNA-Seq analysis of six HIV-associated kidney tumor/adjacent normal tissue samples. We revealed the molecular changes in kidney malignancy among people living with HIV. Our data suggest that certain differentially expressed genes may play key roles in this malignant tumor and may represent novel biomarkers for HIV-associated kidney cancer.

## Materials and Methods

### Patients and tissue specimens

A total of 18 patients presenting at the Shanghai Public Health Clinical Center (Shanghai, China) were diagnosed with HIV-associated kidney cancer from March 2014 to May 2019. The ages of the 18 patients with HIV-associated kidney cancer varied from 24 to 65 years (median, 51 years). The clinicopathological features of the patients included age at diagnosis, sex, cigarette smoking, complications, HAART treatment, CD4^+^ count, surgical approach, pathological type, and TNM stage. Written informed consent was obtained from all patients for the use of tissue samples and clinical records. The study protocol was performed under approval of the Ethics Committee of Shanghai Public Health Clinical Center. All cases were evaluated by two staff pathologists (Dr. J. Xu and Dr. Y. Feng) who were blinded to the clinical outcome and follow-up data.

### RNA purification, whole transcriptome library construction and sequencing

RNA was purified from 18 HIV-associated kidney cancer tumor/adjacent normal tissue samples using TRIzol LS reagent (Invitrogen, Carlsbad, CA, US) and the RNAsimple total RNA kit (Tiangen Biotech Co. Ltd, Beijing, China). The quality of the purified RNA was assessed using an Agilent 2100 bioanalyzer (Agilent Technologies, Waldbronn, DE, US). RNA from HIV-associated kidney tumor tissue samples, including three pairs of tumor/adjacent normal fresh tissue samples, was used for transcriptomic sequencing. Paired-end libraries were synthesized using the TruSeq™ RNA sample preparation kit (Illumina, USA). After poly-A-containing mRNA molecules were purified using poly-T oligo-attached magnetic beads, the mRNA was fragmented into small pieces using divalent cations at 94 °C for 8 min. The cleaved RNA fragments were copied into first strand cDNA using reverse transcriptase and random primers. Then, the second strand of cDNA was synthesized by DNA polymerase I and RNase H. These cDNA fragments then underwent an end repair process with the addition of a single ‘A’ base and the ligation of the adapters. The products were then purified and enriched by PCR to create the final cDNA library. Afterwards, the purified libraries were quantified using a Qubit® 2.0 Fluorometer (Life Technologies, US) and validated using an Agilent 2100 bioanalyzer (Agilent Technologies, US). The library was sequenced on an Illumina NovaSeq 6000 (Illumina, USA). All library construction and sequencing were performed at ShanghaiSinomics Corporation according to the Affymetrix (Santa Clara, CA, US) protocol.

### RNA-Seq data analysis and functional annotation

Fastq files were aligned to hg19 following the best practices for variant calling on RNA-seq as described previously (18). The subsequent gene lists and associated expression values were uploaded to Partek Pro 6.0 software (Partek, MO). When appropriate, the fold change was calculated as the ratio of the mean of gene expression measures of HIV-associated kidney cancer and adjacent noncancer tissue samples (19). To identify potential specific pathways based on changes in gene expression, Gene Ontology (GO) annotation and enrichment were conducted based on the Gene Ontology database (http://www.geneontology.org/). GO-enriched items are shown, and corrected Q values < 0.05 were considered significant. Significantly enriched pathways and biological processes were displayed with R software as described previously (20).

### cBioPortal analysis of The Cancer Genome Atlas data of multiple types of cancer to determine the probability of gene expression alteration and protein– protein interaction (PPI) network analysis

We investigated our candidate genes using cBioPortal analysis of The Cancer Genome Atlas (TCGA) data via cBioPortal (19) and generated the probability of differentially expressed genes (DEGs) for four different types of cancers: kidney renal clear cell carcinoma (TCGA, Nature 2013, n = 392; TCGA, Provisional, n = 446), breast invasive carcinoma (TCGA, Nature 2012, n = 463), bladder urothelial carcinoma (TCGA, Nature 2014, n = 125), and lung adenocarcinoma (TCGA, Nature 2014, n = 230). Next, to explore the interaction of DE-mRNAs of HIV-associated kidney cancer, DE-mRNAs lists were first imported into the STRING database (https://string-db.org), and a protein–protein interaction (PPI) network was generated. When constructing the PPI network, protein–protein interaction pairs with medium confidence of interaction score > 0.9 were exported.

### Quantitative real-time polymerase chain reaction analysis (qRT–PCR)

Total RNA (1 μg) was reverse transcribed to cDNA using the PrimeScriptTM RT reagent kit with gDNA Eraser (Cat^#^: RR047A, TaKaRa Bio, Shiga, Japan). qRT–PCR was performed using a LightCycler 480 II instrument (Roche Molecular Systems, Inc.) and TB Green Premix Ex Taq II (Tli RNase H Plus) (Cat^#^: RR820A, TaKaRa Bio, Shiga, Japan). qPCR results were analyzed using LightCycler^®^ 480 Software Version 1.5 (Roche Molecular Systems, Inc.) and normalized using 18S ribosomal RNA. Eighteen pairs of HIV-associated kidney cancer and adjacent noncancer tissue samples were used for qRT–PCR analysis. The specific primers of the mRNAs are presented in Table S1.

### Immunohistochemical (IHC) staining

IHC staining for KISS1R and CA9 was performed using 18 formalin-fixed, paraffin-embedded tissue samples from the Shanghai Public Health Clinical Center, Fudan University. Paraffin-embedded tissues were dewaxed in xylene, rehydrated by serial concentrations of ethanol, rinsed in phosphate-buffered saline (PBS) and then treated with 3% H_2_O_2_. After being heated at 60 °C overnight, the sections were incubated with 10% normal goat serum at room temperature for 10 min. This step was followed by a PBS wash and incubation with anti-KISS1R (Cat^#^: YT2480, Immunoway Bio., Plano, TX, USA) or anti-CAIX (Cat^#^: YM3076, Immunoway Bio., Plano, TX, USA) antibodies for 12 h at 4 °C followed by horseradish peroxidase (HRP)-conjugated goat anti-rabbit IgG (1/500; Invitrogen, Thermo Fisher Scientific, Inc.). After a PBS wash, the sections were developed in DAB substrate. The sections were then counterstained in hematoxylin for 2 min and dehydrated in ethanol and xylene before being mounted. Eighteen paraffin-embedded tissues were retrieved for IHC analysis. Staining patterns, including cell distribution (membrane, cytoplasmic, and nuclear pattern) an d extent and intensity of staining, were evaluated independently for each specimen by two investigators. For CALX and KISS1R, a 0-4 scale was used for evaluation as follows: negative, focal (< 10%), regional (10–50%), and diffuse (> 50%). The tumors presenting regional and diffuse staining were considered positive, and tumors with negative or patchy staining were categorized as negative.

### Statistical analysis

Data analyses were performed using SPSS statistical package 17.0 (SPSS Inc., Chicago, IL, USA). Statistical values are presented as the means ± standard deviations. Student’s *t*-test was used to assess differences between groups. Univariate analysis was performed using the Kaplan–Meier estimator method and a log-rank test. The median survival time was calculated using SPSS. p < 0.05 was considered to indicate a statistically significant difference.

## Results

### Baseline characteristics

Of the 4027 HIV-positive patients managed by the Shanghai Public Health Clinical Center between March 2014 and June 2019, 18 patients with a diagnosis of RCC were identified, as shown in Table 1, which summarizes the baseline characteristics of the patients with RCC. Among the 18 patients with HIV-associated kidney cancer enrolled in the study, the age of patients with kidney cancer ranged from 24-65 years, and the average age was 51.72 ± 12.15 years. Fifteen of the eighteen patients were on antiretroviral therapy (ART) at the time of RCC diagnosis. Four patients with HIV-associated kidney cancer had histories of comorbidities, such as syphilis or tuberculosis. A total of eight patients were identified with stage I - II kidney cancer, and ten patients were identified with stage III - IV kidney cancer, including thirteen patients with clear cell renal cell carcinoma (ccRcc), three patients with papillary renal cell carcinoma (PRCC), one patient with nephroblastoma, one patient with squamous cell carcinoma (SCC) and one patient with perivascular epithelioid cell tumor (PEComa). Univariate analysis by SPSS showed that overall survival (OS) did not differ by age, sex, smoking, HAART treatment, CD4^+^ count, pathological type, or TNM stage among these patients. The median OS duration of the 18 patients was 18.50 months. However, significant differences in survival outcome were noted between patients without syphilis/hypertension/coronary heart disease/diabetes/hepatitis (30.92 ± 19.53 months) and patients with complications (8.40 ± 3.29 months) (p = 0.023). Specifically, patients with complications were associated with decreased OS duration, an open radical surgical approach (31.08 ± 19.31 months) and an aparoscopic radical surgical approach (8.00 ± 3.74 months) (p = 0.019) (Table 1).

**Table 1.**
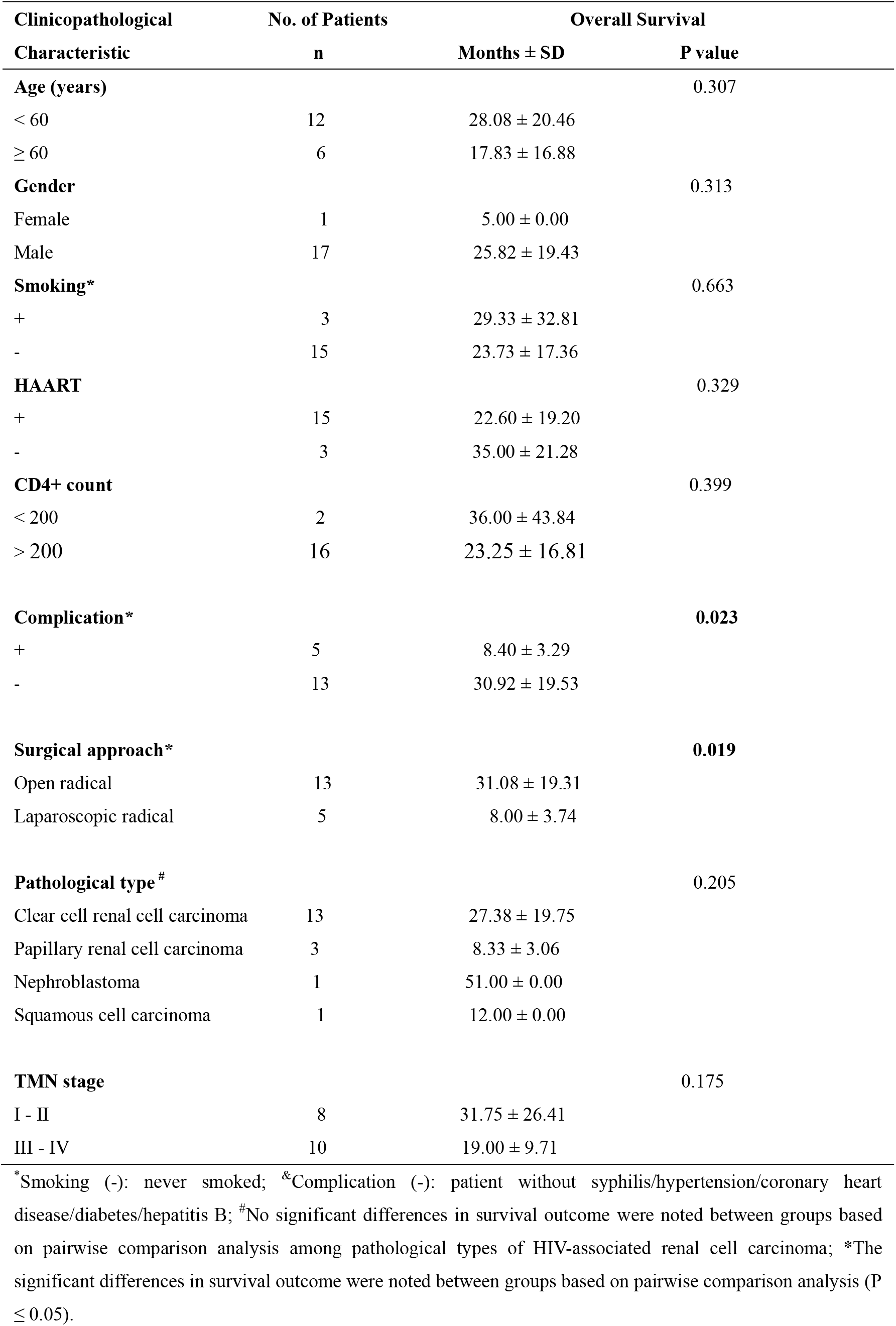
Clinicopathological characteristics of 18 patients with HIV-associated kidney cancer.

### Analysis of RNA-Seq data and transcriptomic profiles in HIV-associated kidney cancer

During the amplification step of sequence generation, the Illumina NovaSeq 6000 produces clusters of identical sequence fragments. The number of these clusters is reported, and the percentage of sequence fragments that passes quality filtering by the Illumina image analysis software is also reported. Across all 6 samples, the total number of reads produced for each sample ranged from 35,739,250 to 45,477,550 with a median of 39,500,005, and 97.61-98.50% of reads were aligned to the reference genome (Table S2). No significant difference in the number of reads from adjacent normal tissue and HIV-associated kidney tumor tissue was noted (p = 0.286). RNA sequencing analysis identified 2460 protein-coding transcripts that exhibited a greater than a 2.0-fold change in expression level (p ≤ 0.05) in the HIV-associated kidney cancer and adjacent normal tissue groups. These DEGs were shown in a volcano plot in Fig. 1A. The heatmap shows that clusters of genes share similar expression patterns at the transcript levels of these DEGs (Fig. 1B) in HIV-associated kidney cancer. Moreover, we identified 555 DEGs (Table 2) with a 5.0-fold change in expression level (Q value ≤ 0.05) in HIV-associated kidney cancer. Furthermore, using TCGA data of multiple cancer types for analysis of 555 DEGs in HIV-associated kidney cancer, we identified that nine DEGs (such as SLC34A1, STC2, FAM153CP, CDHR2, ACOX2, PCDHB1, CHL1, SLC36A2, and FOXI1) had a high incidence of genetic alterations in kidney renal clear cell carcinoma (TCGA-nature 2013, 5.1% - 8.4%; TCGA-provisional, 7.4% - 16.8%) with an incidence cutoff value of ≥ 5.0% in the two kidney cancer TCGA lists that included 838 patients. These genes were also significantly frequently altered across cancer types, including breast invasive carcinoma (1.1% −7.8%), bladder urothelial carcinoma (2.4% - 12.0%), and lung adenocarcinoma (3.0% - 10.9%) (Table 3). Our analysis results suggested that alterations in these nine candidate genes interacted in at least a subset of tumors. A strong tendency of co-occurrences was noted for genetic alterations in these DEGs between ACOX2 and CHL1; SLC36A2 and FOXI1; PCDHB1 and SL36A2 and FOXI1; STC2 and CDHR2, FAM153CP, FOXI1, SLC36A2, and PCDHB1; SLC34A1 and CDHR2, FAM153CP, STC2, FOXI1, SLC36A2, and PCDHB1; FAM153CP and CDHR2, FOXI1, SLC36A2, and PCDHB1; CDHR2 and FOXI1, SLC36A2, and PCDHB1 (p < 0.001) in Table S3. Considering the regulatory role of these nine candidate genes, the underlying mechanisms and cellular consequences of these interactions could be critical for understanding HIV-associated kidney cancer pathology.

**Figure 1:**
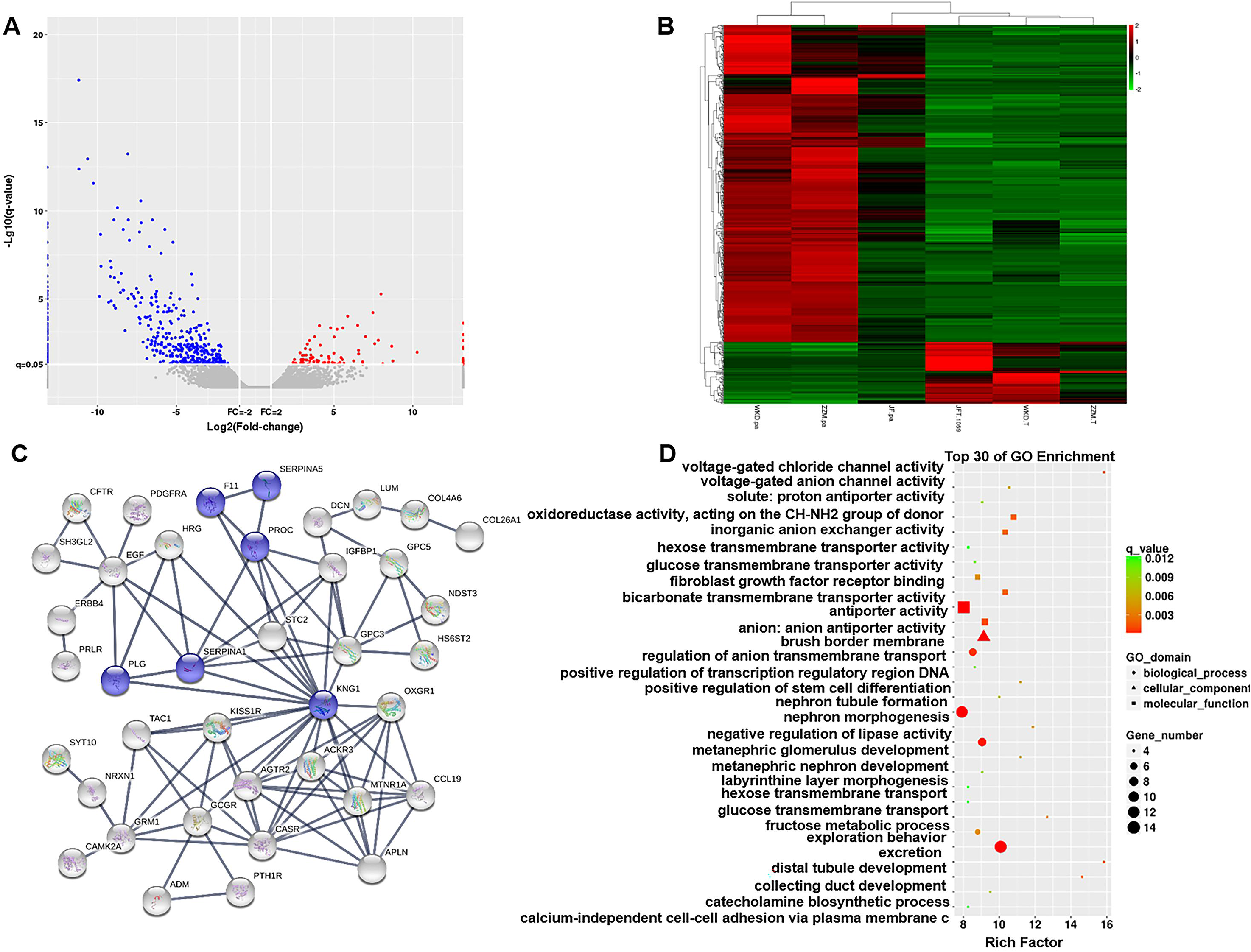
Analysis of significantly expressed genes in HIV-associated kidney cancer, which are shown in a volcano plot (A) and based on hierarchical clustering (C). Protein–protein interaction analysis (A) and functional enrichment analyses of mRNAs (C) functional network analysis in HIV-associated kidney cancer. Upregulated DEGs are shown in red, and downregulated DEGs are shown in green. Compared with adjacent normal tissue groups, significantly highly expressed genes in HIV-associated kidney cancer were defined as genes with a fold change ≥ 2.0, P ≤ 0.05, shown in red; significantly less expressed genes are shown in green/blue with a fold change ≤ 0.5, P value ≤ 0.05; genes that did not meet the criteria are shown in gray.

**Table 2.** The partial differentially expressed genes in HIV-associated kidney cancer.

| gene name | description | FC | P value |
| --- | --- | --- | --- |
| AC079466.1 | - | 1230.92 | 0.010 |
| KISS1R | KISS1 receptor | 403.11 | 0.005 |
| CAIX | carbonic anhydrase 9 | 251.39 | 0.000 |
| TEX15 | testis expressed 15, meiosis and synapsis associated | 223.10 | 0.002 |
| IGFBP1 | insulin like growth factor binding protein 1 | 190.32 | 0.003 |
| EGFR-AS1 | EGFR antisense RNA 1 | 176.81 | 0.000 |
| NPTX2 | neuronal pentraxin 2 | 127.33 | 0.001 |
| AF064858.1 | - | 93.96 | 0.008 |
| AL928654.4 | - | 89.82 | 0.000 |
| GRIK3 | glutamate ionotropic receptor kainate type subunit 3 | 58.26 | 0.000 |
| ANGPTL4 | angiopoietin like 4 | 46.88 | 0.000 |
| SLC17A4 | solute carrier family 17 member 4 | 36.50 | 0.001 |
| PGF | placental growth factor | 32.19 | 0.001 |
| HILPDA | hypoxia inducible lipid droplet associated | 27.53 | 0.000 |
| GAS6-AS1 | GAS6 antisense RNA 1 | 17.43 | 0.004 |
| SLC12A3 | solute carrier family 12 member 3 | -489.44 | 0.000 |
| CYP4F2 | cytochrome P450 family 4 subfamily F member 2 | -494.43 | 0.000 |
| SLC12A1 | solute carrier family 12 member 1 | -501.39 | 0.000 |
| SLC36A2 | solute carrier family 36 member 2 | -564.26 | 0.000 |
| NPHS1 | NPHS1, nephrin | -566.76 | 0.000 |
| SLC4A1 | solute carrier family 4 member 1 (Diego blood group) | -585.42 | 0.000 |
| PLA2G4F | phospholipase A2 group IVF | -588.01 | 0.000 |
| CALB1 | calbindin 1 | -631.08 | 0.000 |
| SLC22A8 | solute carrier family 22 member 8 | -883.25 | 0.000 |
| SLC13A2 | solute carrier family 13 member 2 | -902.43 | 0.000 |
| NPHS2 | NPHS2, podocin | -945.21 | 0.000 |
| CLDN8 | claudin 8 | -1228.85 | 0.000 |
| TMEM213 | transmembrane protein 213 | -1583.83 | 0.000 |
| AQP2 | aquaporin 2 | -2324.43 | 0.000 |
| UMOD | uromodulin | -2337.55 | 0.000 |

**Table 3.** The Cancer Genome Atlas consortium data on the incidence of genetic

| GENE_SY<br>MBOL | Kidney Renal Clear Cell<br>Carcinoma (n = 392) |  | Kidney Renal Clear Cell<br>Carcinoma (n = 446) |  | Breast Invasive<br>Cancer (n = 463) |  | Bladder Urothelial<br>Carcinoma (n =<br>125) |  | Lung Adenocarcinoma<br>(n = 230) |  |
| --- | --- | --- | --- | --- | --- | --- | --- | --- | --- | --- |
|  | n | % | n | % | n | % | n | % | n | % |
| SLC34A1 | 33 | **8.4%** | 74 | **16.6%** | 23 | **5.0%** | 11 | **8.8%** | 23 | **10%** |
| STC2 | 27 | **6.9%** | 75 | **16.8%** | 9 | 1.9% | 7 | **5.6%** | 17 | **7.4%** |
| FAM153C<br>P | 23 | **5.9%** | 69 | **15.5%** | 5 | 1.1% | 8 | **6.4%** | 13 | **5.7%** |
| CDHR2 | 23 | **5.9%** | 74 | **16.6%** | 5 | 1.1% | 9 | **7.2%** | 20 | **8.7%** |
| ACOX2 | 22 | **5.6%** | 33 | **7.4%** | 19 | 4.1% | 10 | **8.0%** | 7 | 3.0% |
| PCDHB1 | 21 | **5.4%** | 66 | **14.8%** | 29 | **6.3%** | 7 | **5.6%** | 16 | **7.0%** |
| CHL1 | 20 | **5.1%** | 56 | **12.6%** | 30 | **6.5%** | 15 | **12.0%** | 25 | **10.9%** |
| SLC36A2 | 20 | **5.1%** | 65 | **14.6%** | 36 | **7.8%** | 3 | 2.4% | 8 | 3.5% |
| FOXI1 | 20 | **5.1%** | 70 | **15.7%** | 25 | **5.4%** | 5 | 4.0% | 11 | 4.8% |

alterations of DEGs in HIV-associated kidney cancer based on cancer type.
<sup>a</sup>Genetic alterations include mutations and/or CNV. <sup>b</sup>Cancer Genome Atlas Research Network, Nature, 2014 [15].

### Protein–protein interaction (PPI) and functional pathway analysis of HIV-associated kidney cancer

To further elucidate the regulatory and interaction relationships, 555 DEGs were imported into the STRING database. The STRING database identified a network of 462 mRNAs and 184 edges (interaction relationship). The 6 mRNAs (F11, SERPINA5, SERPINA1, PLG, KNG1, and PROC) in the network were reported to regulate the complement and coagulation cascades (Fig. 1C). Subsequently, GO analysis of these 555 differentially expressed mRNAs was performed to determine the top 30 of GO enrichment categories of these differential mRNAs in HIV-associated kidney cancer. Several common GO terms were identified for the differentially expressed mRNAs (Fig. 1D) in HIV-associated kidney cancer, including voltage-gated chloride channel activity, distal tubule development, collecting duct development, fructose metabolic processes, and negative regulation of lipase activity.

### Quantitative real-time PCR validation of DEGs in HIV-associated kidney cancer

Real-time PCR (qRT–PCR) analyses of KISS1R, CAIX, NPTX2, TMEM213, and UMOD mRNA expression in 18 pairs of HIV-associated kidney cancer and adjacent noncancer tissue samples revealed that 18 of 18 (100.0%) tumors had increased NPTX2 mRNA (110.90-fold) (p = 0.027), 13 of 18 (72.2%) tumors had increased KISS1R mRNA (92.08-fold) (p = 0.020), and 14 of 18 (77.8%) had increased CAIX mRNA (225.84-fold) (p = 0.012) expression in HIV-associated kidney cancer (Fig. 2A-2C). In addition, we also found that 15 of 18 (78.6%) tumors had decreased UMOD mRNA (2308.15-fold) (p = 0.040), and 14 of 18 (77.8%) had decreased TMEM213 mRNA (405.95-fold) (p = 0.037) expression in HIV-associated kidney cancer (Fig. 2D and 2E). To test whether these DEGs were unique to HIV-associated kidney cancer, we analyzed their expression levels in 34 patients with kidney cancer. Thirty-two of 34 (94.1%) tumors had increased NPTX2 mRNA (253.22-fold) (p = 0.052), 30 of 34 (88.2%) tumors had increased KISS1R mRNA (94.00-fold) (p = 0.167), and 29 of 34 (85.3%) had increased CAIX mRNA (855.56-fold) (p = 0.058) expression in kidney cancer (Fig. 2F–3H). We also found that 33 of 34 (97.1%) tumors had decreased UMOD mRNA (789.50-fold) (p < 0.001), and 31 of 34 (91.2%) had decreased TMEM213 mRNA (893.78-fold) (p = 0.002) in kidney cancer (Fig. 2I and 2J). Our results indicated that HIV-positive renal cell carcinoma (RCC) patients were similar to the general population in terms of pathology.

**Figure 2:**
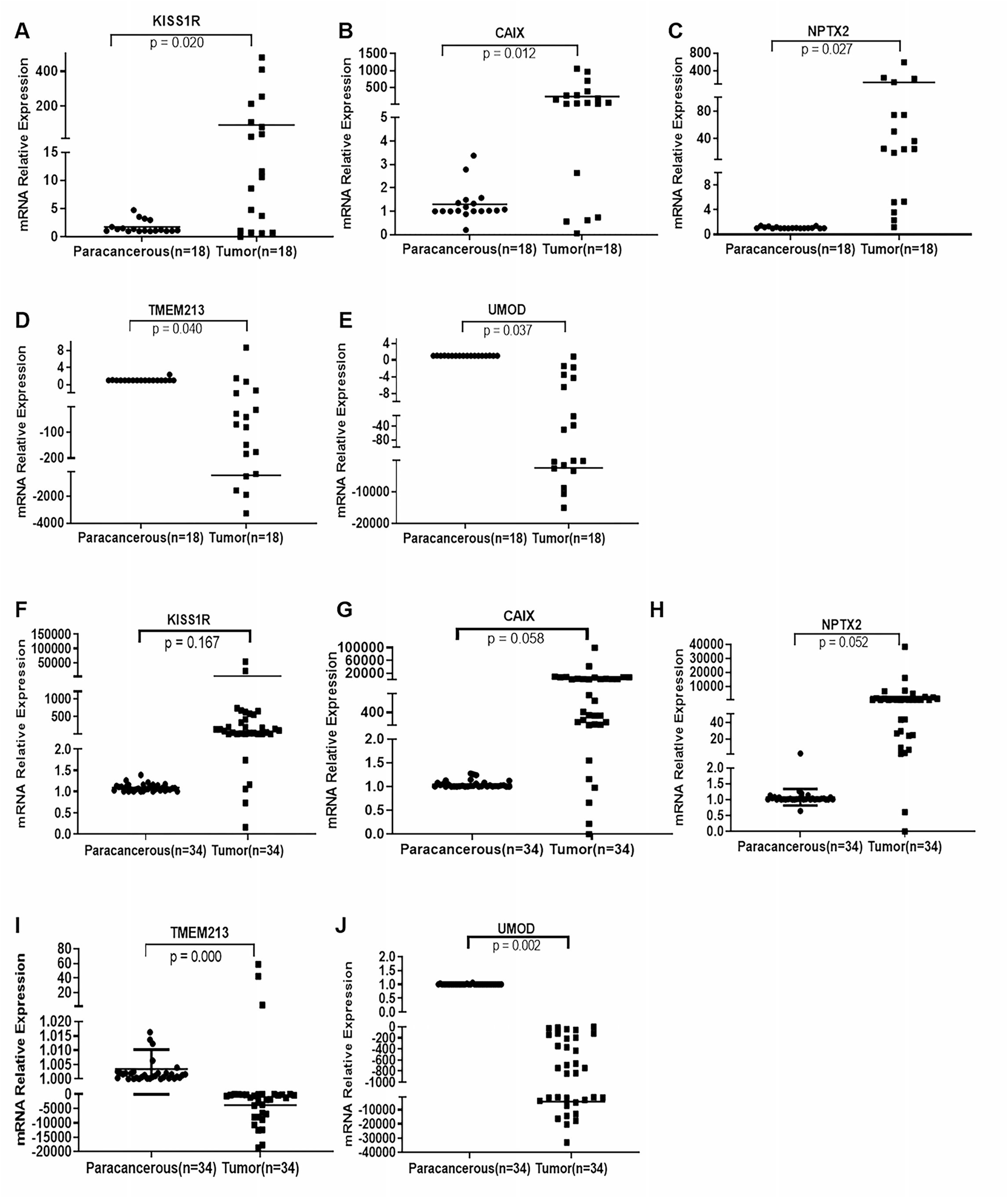
Real-time PCR (qRT–PCR) analysis of KISS1R, CAIX, NPTX2, TMEM213, and UMOD mRNA expression in 18 pairs of HIV-associated kidney cancer and adjacent noncancer tissue samples; 34 pairs of kidney cancer and adjacent noncancer tissue samples (Cont: adjacent noncancer tissue; Ca: kidney tumor tissue). A-E) HIV-associated kidney tumor and adjacent noncancer tissue samples; F-J) kidney tumor and adjacent noncancer tissue samples; A, F) KISS1R; B, G) CAIX; C, H) NPTX2; D, I) TMEM213; E, J) UMOD.

### IHC staining analysis of KISS1R and CAIX expression in HIV-associated kidney cancer

To investigate the expression of KISS1R and CAIX proteins in benign and malignant kidney tissue, we performed HE and IHC staining of 18 HIV-associated kidney tumor sample samples (Fig. 4). In total, KISS1R- and CAIX-specific staining was clearly observed in the cytoplasm and membrane of primary kidney cancer cells. We observed positive staining of KISS1R and CAIX in HIV-associated kidney ccRcc, squamous cell carcinoma (SCC), and papillary renal cell carcinoma (PRCC) (Fig. 3). HE staining of HIV-associated kidney tumors (HE × 400) showed that the tumor cells of grade I ccRc in Fig. 3A were large with small nuclei, transparent cytoplasm, inconspicuous nucleoli and no mitosis (red arrow refers to ccRc). The cells of grade II ccRC in Fig. 3D were also large with obvious nucleoli and no mitotic figures. Most of the cells had transparent cytoplasm, and some of the cells were eosinophilic small granular cells where the blood vessels were visible in the interstitium (red arrows indicate ccRcc). The tumor cells of grade Ⅰ-Ⅱ SCC in Fig. 3G were also large, and most of the cells were eosinophilic cells with inconspicuous nucleoli and occasional mitotic figures. These cells were arranged in clusters in the stroma (red arrow refers to SCC) (Fig. 3G). The cells of grade I PRCC in Fig. 3J were medium, papillary shape, cytoplasmic eosinophilia with inconspicuous nucleoli and no mitotic figures (red arrows refer to PRCC tumor cells). Staining of HIV-associated kidney tumors was variable, ranging from cytoplasmic to membranous with poorly differentiated tumor tissues of cRcc (grade II) showing higher expression of KISS1R (Fig. 3E) and CAIX (Fig. 3F) (IHC × 400) compared to the expression of KISS1R (Fig. 3B) and CAIX in grade I ccRc (Fig. 3C). We found that KISS1R was almost negatively expressed in grade I ccRc tumor cells (Fig. 3B) but was widely expressed in the cytoplasm of grade II ccRc and PRCC tumor cells, partly in their nuclear membrane (Fig. 3E and 3K). KISS1R also strongly expressed the nuclear membrane of SCC tumor cells, and only some of them were expressed in the cytoplasm (Fig. 3H). We also found that CAIX was expressed in the cell membrane of grade I ccRc (Fig. 3C), and it was also strongly positively expressed in the nuclear membrane and cytoplasm of grade II ccRc tumor cells (Fig. 3F). Although CAⅨ was expressed in the cytoplasm of PRCC tumor cells (Fig. 3L), weak positive CAⅨ expression was verified in the cytoplasm of SCC tumor cells (Fig. 3I) (IHC × 400). A total of 16 of 18 (88.9%) malignant cases showed positive staining for KISS1R and CAIX. All these results indicated that the expression levels of KISS1R and CAIX are specifically increased in HIV-associated kidney cancer.

**Figure 3:**
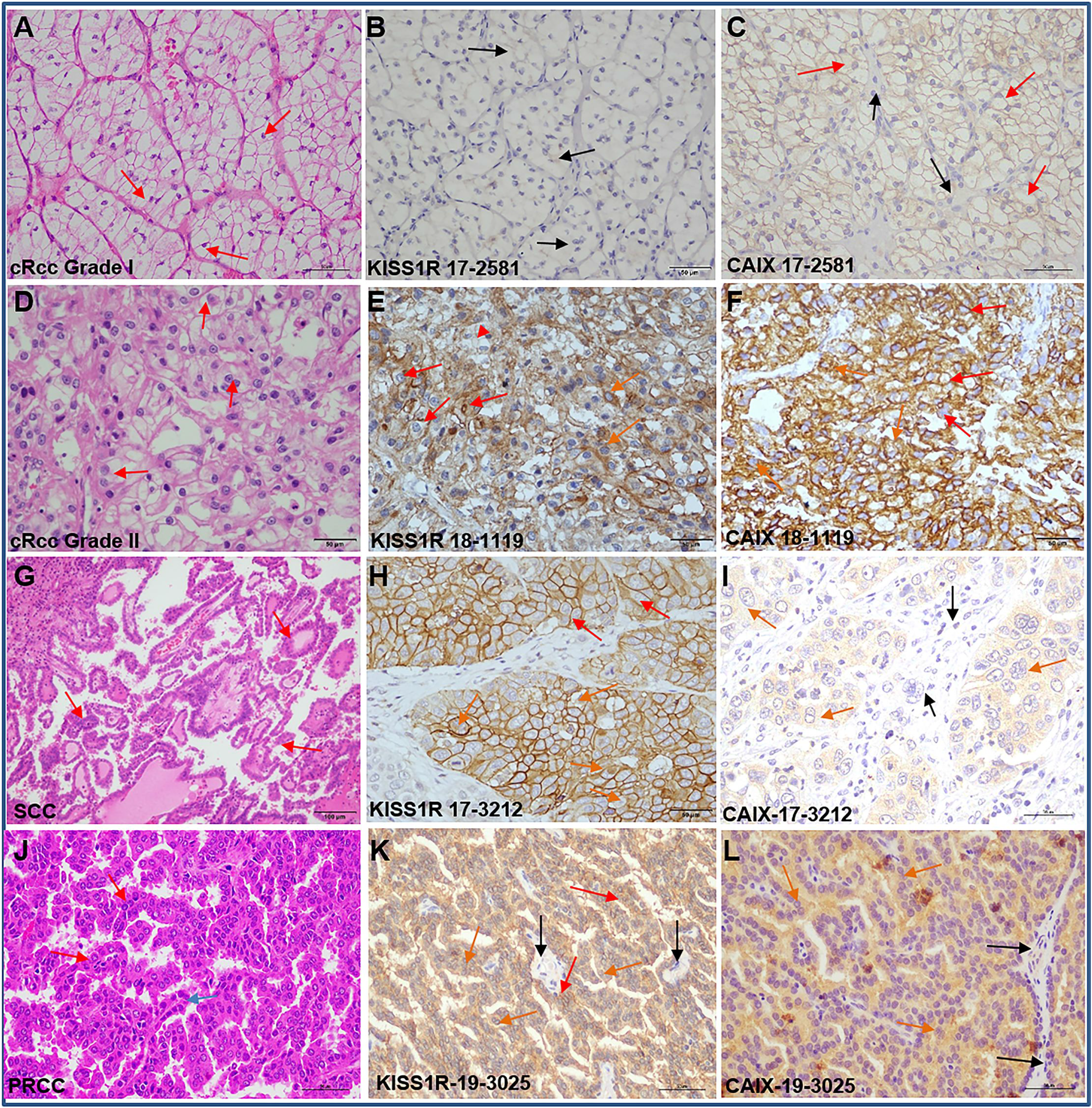
Immunohistochemical analysis of KISS1R and CAIX protein expression in HIV-associated kidney tumor tissue samples stained with anti-KISS1R (B, E, H and K), anti-CAIX antibodies (C, F, I and L), and HE staining of cRcc tumor tissue samples (A, D, G and J). Staining of variably differentiated kidney tumor samples with an anti-KISS1R antibody (B, E, H and K) and an anti-CAIX antibody (C, F, I and L) (red arrows refer to KISS1R/CAⅨ expressed in the cell membrane or nuclear membrane, orange arrows refer to KISS1R/CAⅨ expressed in the cell cytoplasm, and black arrows indicate negative KISS1R/CAIX expression; IHC × 400).

## Discussion

RNA-seq of tumor specimens can help to shed light on the possible pathogenicity of variants of unknown significance because these data provide direct insight into transcriptional alterations and thus improve diagnostic rates(21). Although we previously identified 758 differentially expressed genes in HIV-associated lung cancer and reported that SIX1 and TFAP2A were overexpressed in HIV-associated lung cancer and associated with poorly differentiated tumors (19), known molecular changes in kidney malignancy among people living with HIV are limited. This information is important to investigate the molecular mechanisms of HIV-associated kidney cancer. In this study, we first analyzed the baseline characteristics of 18 patients with HIV-associated kidney cancer enrolled in our center. The baseline characteristics of these patients are similar to those of 7 patients with HIV-associated renal cell carcinoma enrolled in an Australian statewide HIV center. The median age at RCC diagnosis was 56 years. In addition, six of the seven patients were on ART at RCC diagnosis, and five had virological suppression. Two patients had metastatic RCC at diagnosis, and the other five patients with clinically localized RCC had radical/partial nephrectomies. The median OS duration of the four patients who died of metastatic RCC was 9 months, and the other three patients were alive at a median follow-up of 16 months (22). These studies suggested that effective biomarkers should be used for early staging of cancer, and personalization of therapy at the time of diagnosis should be offered, which may improve HIV-associated kidney cancer patient care. Thus, in our initial biomarker identification stage, transcriptomic sequencing was performed to assay the differentially expressed genes in HIV-associated kidney cancer tumor tissue samples, including three pairs of tumor/adjacent normal fresh tissue samples. We found 2279 DElncRNA transcripts and 2460 DEGs/mRNAs in HIV-associated kidney cancer compared to the adjacent normal tissue groups. Among them, nine DEGs (SLC34A1, STC2, FAM153CP, CDHR2, ACOX2, PCDHB1, CHL1, SLC36A2, and FOXI1) also had a high incidence of genetic alterations in kidney renal clear cell carcinoma in the two kidney cancer TCGA lists of 838 patients.

Next, we identified that several pathways from both differentially expressed mRNAs in HIV-associated kidney cancer, including the structure and function of key transporters, tubule and duct development, fructose metabolic processes, and regulation of lipase activity, which may be altered in HIV-associated kidney cancer. Coding variants in apolipoprotein L1 (APOL1), which has broad innate immune functions and can restrict HIV replication in vitro, are strongly associated with HIV-associated nephropathy in persons with untreated HIV infection (23). Strong associations in genome-wide association studies were noted between chronic kidney disease and UMOD promoter variants, which influenced urinary uromodulin levels (24). Uromodulin encoded by UMOD can be secreted and passively excreted in urine (25). Several TMEM family members were deregulated in ccRcc tumors as components of cellular membranes (such as mitochondrial membranes, ER, lysosomes, and Golgi apparatus), suggesting their importance in the pathogenesis of ccRcc with the involvement of ER proteins (26). Our qRT–PCR analyses also validated that TMEM213 and UMOD were repressed in HIV-associated kidney cancer (Fig. 2A-2E), and KISS1R, CAIX, or NPTX2 were significantly upregulated in HIV-associated kidney cancer (p < 0.05) (Fig. 2A-2C). Positive staining for KISS1R and CAIX was further observed in HIV-associated kidney ccRcc (Fig. 3C, 3E and 3F), squamous cell carcinoma (Fig. 3H and 3I), and papillary renal cell carcinoma (Fig. 3K and 3L). The differentially expressed genes CAIX, NPTX2 and UMOD were also found in the formalin-fixed, paraffin-embedded (FFPE) and RNAlater® datasets and confirmed by immunohistochemistry (27). More importantly, we found higher expression of KISS1R (Fig. 3E) and CAIX in the tumor tissues of grade II cRcc (Fig. 3E and 3F) compared to KISS1R and CAIX expression in grade I ccRc (Fig. 3B and 3C), which demonstrated that increased KISS1R and CAIX expression was closely linked to poor clinical prognosis of cancer patients. The cytokeratin (CK) AE1/AE3 and vimentin immunoexpression were also analyzed in 26 CCRCCs. CK AE1/AE3 immunoexpression was associated with low-grade and early-stage lesions. Vimentin immunoexpression was also associated with high-grade and advanced lesions (28). In clear cell RCC, 89.2% of cancers showed strong positivity for CAIX staining. The frequency and intensity of CAIX staining varied between different RCC tumor types, and CAIX staining was strong in 61.2% of 1677 tumor tissue samples (29). On the other hand, the KISS1 protein can be rapidly processed in serum into smaller but biologically active peptides called kisspeptins through the G-protein coupled receptor KISS1R. KISS1, kisspeptins and KISS1R might regulate the development and progression of several cancers, including melanoma and pancreatic, colorectal, bladder, and ovarian cancer (30). We also analyzed KISS1R, CAIX, NPTX2, TMEM213 and UMOD expression in 34 pairs of kidney tumor compared with adjacent noncancer tissue samples from the general kidney cancer population and revealed that the expression levels of KISS1R (p = 0.167), CAIX (p = 0.058), and NPTX2 (p = 0.052) were not significantly increased in kidney tumor tissue from the general kidney cancer population. These results implied that KISS1R might represent a differentially expressed gene that is specific for HIV-infected kidney cancer.

Finally, the limitations of the current study should be noted. Our current sample size of patients with non-HIV kidney cancer or HIV-associated kidney cancer is small. The limitations of this study are that the lack of tissue for comparison limits the ability to determine the specificity of differential gene expression for HIV-infected kidney cancer. It would be interesting to find other differentially expressed genes or pathways in HIV-infected kidney cancer that are specific to HIV. Against this background, there is an opportunity to develop novel gene signatures for HIV-associated cancer that are useful for the early detection of HIV-associated kidney cancer. The current study is notable because the information reported herein may be useful for early detection as well as tracking disease progression and recurrence, which would help kidney cancer screening in HIV-infected populations and guide treatment of this disease.

## Acknowledgments

This research was supported by a grant from the Science and Technology Commission of Shanghai (20Y11900700), a grant from Shanghai Natural Science Foundation (20ZR1470500), a grant from the Special Research Fund of Youan Medical Alliance for the Liver and Infectious Diseases (LM202020), and a grant (KY-GW-2020-09) (X. He) from Shanghai Public Health Clinical Center, Shanghai, China.

## Abbreviations

AIDS: Acquired immunodeficiency syndrome
ART: Antiretroviral Therapy
ccRcc: clear cell renal cell carcinoma
FFPE, DEGs: differentially expressed genes
Formalin-fixed: paraffin-embedded
HAART: Highly active antiretroviral therapy
HIV: Human immunodeficiency virus
IHC: Immunohistochemical
IPA: Ingenuity Pathway Analysis
NADMs: non-AIDS–defining malignancies
OS: Overall survival
PPI: protein–protein interaction
PEComa: perivascular epithelioid cell tumor
PLWH: People living with HIV
PRCC: Papillary renal cell carcinoma
qRT–PCR: Quantitative real-time PCR
RCC: renal cell carcinoma
SCC: squamous cell carcinoma
TCGA: The Cancer Genome Atlas

